# ANTIGEN-SPECIFIC CD8+ T-CELLS ARE INVOLVED IN HUMAN CAROTID ATHEROSCLEROTIC PLAQUES DESTABILIZATION

**DOI:** 10.1101/2025.09.15.676431

**Authors:** Dalgisio Lecis, Manuel Scimeca, Fabio Massimo Oddi, Alessandro Mauriello, Arnaldo Ippoliti, Francesco Buccisano, Maria Antonietta Irno Consalvo, Gianluca Massaro, Daniela Benedetto, Paul C Dimayuga, Kuang-Yuh Chyu, Prediman Krishan Shah, Giuseppe Sangiorgi

## Abstract

Immunization of ApoE^-/-^ mice expressing human HLA-A 02:01 with p210, an apoB100-derived peptide, reduces atherosclerotic plaque development by inducing a p210-specific CD8+ T cell population. Studying Class-I MHC/CD8+ T cell signaling offers a promising approach to understanding the mechanism behind the athero-protective effects of p210 immunization. We aimed to identify a p210-specific CD8+ T cell population in human carotid atherosclerotic plaques from blood-positive HLA-A 02:01 patients undergoing surgical carotid endarterectomy (CEA). The study included 22 consecutive patients who were HLA-A 02:01 (+) out of 49 enrolled (reflecting an estimated prevalence of about 30% HLA-A02:01(+) in the Caucasian population). Immunohistochemistry staining used a PE-marked A 02:01–KTTKQSFDL Pentamer on fixed endarterectomy plaques. Both HLA-A 02:01 (+) and (-) patient plaques were used, with the latter serving as an internal negative control. Presence of pentamer (+) CD8 T cells indicated a p210-specific CD8+ T cell population. Patients positive for HLA-A 02:01 showed an average of 3.40 ± 2.17 × 10^3 HPF p210-specific CD8+ T cells (61.80%, 3% of total CD3+) in the shoulders of atherosclerotic plaques post-CEA, significantly higher than in controls (p < 0.0001). The proportion of p210-specific CD8+ T cells was lower in plaques displaying morphological features of instability. This study, for the first time, identifies a p210-specific CD8+ T cell population in human carotid atherosclerotic plaques from HLA-A02:01(+) patients, suggesting a role for autoimmunity in atherosclerosis development and supporting the potential efficacy of p210 immunization in HLA-A02:01 (+) individuals to reduce atherosclerosis. The variation in this specific T cell population within human plaques correlates with plaque vulnerability, highlighting p210-specific CD8+ T cells as a potential target for future therapies.

## INTRODUCTION

Atherosclerosis is a chronic inflammatory disease characterized by the buildup of pro-atherogenic substances in the arterial intima layer. Both innate and adaptive immune responses contribute to promoting the inflammatory process of atherosclerosis, likely to remove potentially harmful agents from the arterial wall. Some studies have identified various infectious microorganisms in the arterial wall that may serve as potential antigens, triggering inflammation. However, recent evidence indicates that the primary targets of these immune responses are modified endogenous structures.

Studies have shown that in hypercholesterolemic mice lacking specific components of the innate or adaptive immune system, the development of atherosclerosis is reduced, highlighting the role of immunity in this complex disease. Additionally, research on genetic differences in the HLA (human leukocyte antigen) complex, which is crucial for presenting antigens to lymphocytes, has identified various variants associated with a higher risk of cardiovascular disease^6^. This strong connection between the immune system and atherosclerosis has led scientists to view this process, at least partly, as an autoimmune disease, while also recognizing that metabolic and hemodynamic factors act as initial triggers. The role of inflammation and the immune system in atherosclerosis is to clear waste products, such as degradation fragments from oxidized LDL (oxLDL). In recent years, efforts have focused on identifying these waste products, also known as self-antigens, that trigger inflammation and contribute to the development of atherosclerosis.

The most studied and characterized self-antigen in atherosclerosis is oxLDL, which forms through enzymatic attacks on retained LDL in the arterial wall’s intima. This process releases pro-inflammatory phospholipids and lipid peroxides, triggering a rapid inflammatory response. Oxidation causes the fragmentation of apoB-100, the main apoprotein of LDL, and produces various aldehyde and phospholipid adducts on apoB-derived peptides^8^. These peptides can promote the clonal expansion of T cells specific to oxidized LDL, leading to the production of certain autoantibodies.

As the importance of immunity in atherosclerosis has been revealed, it is interesting to clarify whether this knowledge can be applied to develop new treatments for cardiovascular disease. The first evidence that this could be possible came from studies in which hypercholesterolemic rabbits were immunized with oxidized LDL^9,10^. The initial aim of these studies was to test whether activation of immunity to oxidized LDL was associated with a more aggressive progression of disease. However, it was found, surprisingly, that immunization against oxidized LDL actually reduced atherosclerosis in animal models. This observation was subsequently confirmed in several different animal models of atherosclerosis, suggesting the fascinating possibility that a vaccine could be developed for atherosclerosis. However, since oxidized LDL is a complex particle with an antigen composition that is difficult to standardize and may potentially contain harmful antigens, it is not an ideal vaccine component on its own. Over the last few years, considerable efforts have therefore been made to characterize the precise antigens and antigenic epitopes in oxidized LDL that induce athero-protective immunity.^11^

Fredrikson et al. demonstrated that immunizing ApoE-/-mice with peptides derived from apo-B100, the main protein of LDL, reduced the development of atherosclerotic plaques. Using a library of polypeptides covering the entire sequence of apoB-100, they identified over 100 different peptides within apoB-100 that could serve as potential antigens^13^. Further testing showed that peptides like p2, p143, and p210 led to a 40% to 70% reduction in atherosclerosis and decreased plaque inflammation when used in a vaccine formulation in hypercholesterolemic mice. Since the p210-based vaccine produced the most consistent anti-atherosclerotic effects, this peptide was selected as a prototype antigen for vaccine formulations^15,16^.

Chyu et al. demonstrated that immunizing mice with p210 reduced atherosclerotic plaque development, and the protective effect is mediated by a CD8+ T cell population. Then, Dimayuga et al. identified a p210 fragment-specific CD8+ T cell population in ApoE-/-mice fed an atherogenic diet, using a pentamer consisting of HLA class I H2-Kb molecules binding a p210-derived octamer. Subsequent immunization of mice with p210 showed a shift in the immune-dominant epitope. Recently, the potential clinical relevance of immunization with p210 in self-assembling peptide amphiphile micelles (P210-PAM), a novel vaccine formulation, was demonstrated by reducing atherosclerosis in a humanized ApoE-/-mouse model expressing chimeric HLA-A^*^02:01/Kb^19^.

The immune response relies on cell expression of specific HLA molecules. Building on previous research, studying class-I HLA/CD8+ T cell signaling is a promising way to understand the mechanism behind the athero-protective effects of p210 immunization. The allele HLA-A*02:01 is the most common among Caucasians.

In this study, we collected 49 consecutive atherosclerotic plaques from male and female patients who underwent carotid endarterectomy (CEA). A blood sample from each patient was used to identify those with HLA-A*02:01. All HLA-A*02:01+ human plaques were analyzed by fluorescence microscopy after staining with a fluorescently labeled p210/A*02:01 Pentamer and a mouse anti-CD3 antibody to identify a p210-specific CD8+ T cell population. Based on these considerations, this study aimed to investigate the presence of a p210-specific CD8+ T cell population in carotid plaques and its association with plaque instability.

## MATERIALS AND METHODS

The present study was conducted in accordance with the Declaration of Helsinki, the principles of Good Clinical Practice, and GDPR 679/16. The Policlinico Tor Vergata Ethics Committee approved the study (protocol #116.22).

### INCLUSION CRITERIA

Every effort has been made to verify participants’ eligibility before enrollment. Patients who did not meet all the inclusion/exclusion criteria were not enrolled in the study.

Candidates for this study met the following criteria: male or female, capable of understanding and signing a witnessed informed consent form for blood sample collection; age ≥ 18 years; patients with carotid stenosis over 70% and an indication for CEA; and a life expectancy greater than 1 year.

### EXCLUSIONS CRITERIA

Candidates were considered ineligible for enrollment in the study if any of the following conditions applied: active cancer treated with chemotherapy or radiation; patients taking immunosuppressive drugs; pregnant women.

### POPULATION CHARACTERISTICS

The patient’s clinical and laboratory characteristics collected included age, sex, cardiovascular risk factors, BMI, mean LDL and HDL cholesterol levels, and use of cholesterol-lowering medications. Additionally, we examined the presence of symptoms related to significant carotid artery stenosis, such as transient ischemic attack or minor stroke within the six months prior to CEA.

### HLA A*02:01 CHARACTERIZATION

A 10 mL EDTA tube containing blood from patients who underwent CEA was sent to the OPPO laboratory at Policlinico Tor Vergata. A bulk lysis was performed using 1 mL of peripheral blood. The pellet was resuspended in physiological solution, and 100 µL was incubated in a polypropylene tube with the following antibodies: HLA-A*02:01 PEA, CD3 PerCP-Cy5.5, CD4 PE-Cy7, CD8 APC-H7, and CD45 V500 (BD Biosciences). A second tube was prepared with the same antibody combination but without the HLA-A*02:01 antibody, serving as the internal negative control. After a 15-minute incubation in the dark, the samples were analyzed on a BD FACSlyric flow cytometer using BD FACSuite software. Figure (Fig.) 1 displays the gating strategy for identifying HLA-A*02:01+ patients.

**Fig. 1.**
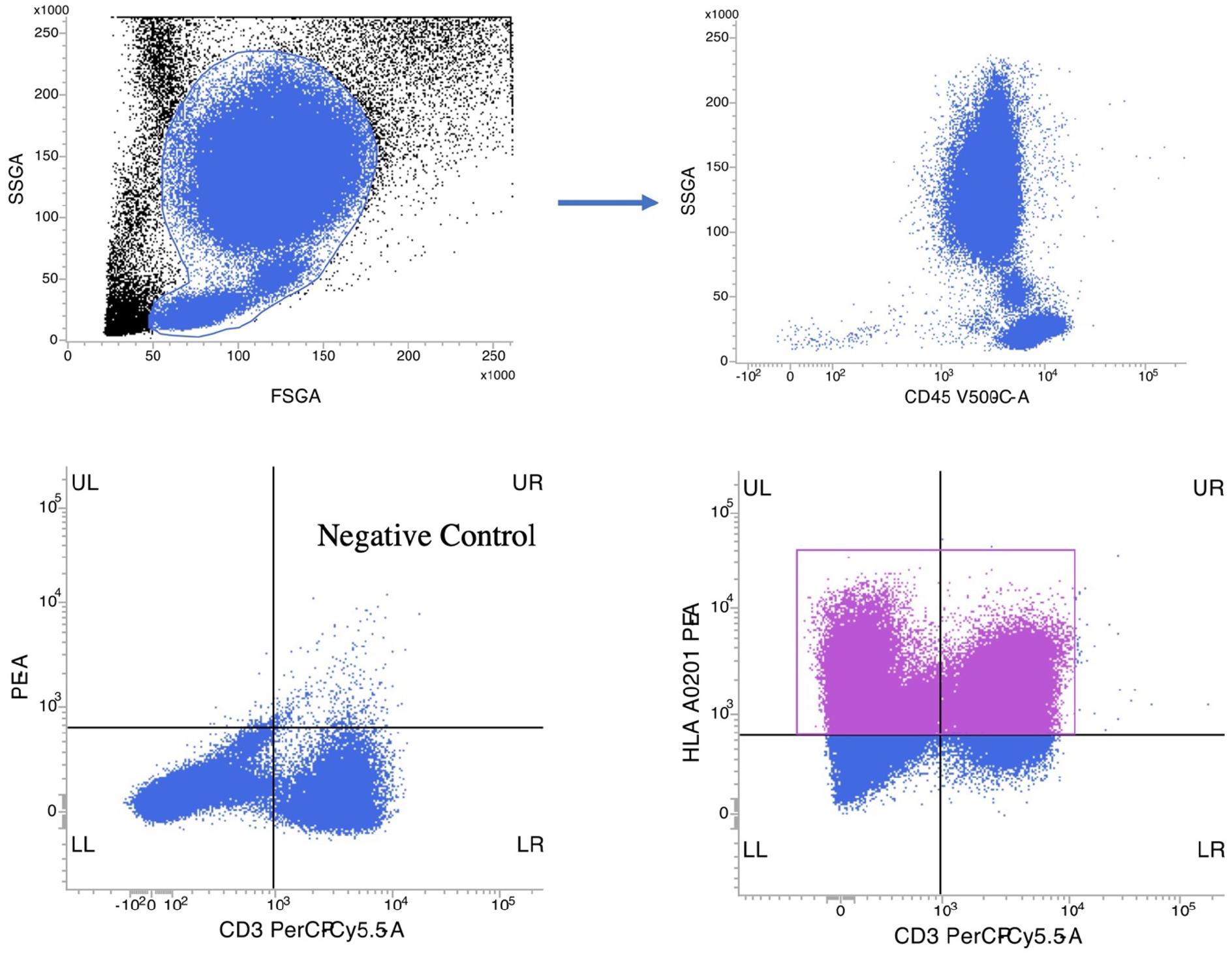
Gating strategy for the identification of HLA-A*02:01 patients.

### ATHEROSCLEROTIC PLAQUES STAINING

The Pentamer staining was performed according to the manufacturer’s instructions, using the HLA-A*02:01 Negative Pentamer loaded with an irrelevant peptide (ProImmune) as the negative control. We also tested the p210/A*02:01 pentamer on a few HLA-A*02:01 patients to confirm its specificity, acting as an internal control.

Specifically, each section was incubated with 20 µL of fluorescently labeled MHC Pentamer (red) for 30 minutes. To achieve co-localization of the pentamer and CD3, the sections were then incubated for 30 minutes with mouse anti-CD3 antibodies (clone CD3-565-L-CE; pre-diluted; Leica, Heidelberg, Germany). A FITC-conjugated anti-mouse secondary antibody was used to detect the primary antibody. For nuclear staining, DAPI incubation was performed on the sections. The quantification of pentamer-CD3 co-positive cells was carried out using a fluorescence microscope, evaluating three high-power fields (HPF) at 63x magnification. Subsequently, to confirm the T cell phenotype, a second staining experiment was conducted using a pentamer (red) and a mouse anti-CD8 antibody detected with a FITC anti-mouse secondary antibody.

### STATISTICS

GraphPad Prism 9.0 software was used for statistical analysis. Data are shown as mean ± SD. The Wilcoxon signed-rank test compared HLA-A*02:01+ plaques stained with the p210/A*02:01 Pentamer and the A*02:01-negative pentamer (controls). To compare the total number of CD3+ T cells and the percentage of p210-specific CD8+ T cells between symptomatic and asymptomatic patients, a Mann-Whitney test was employed. A p-value of less than 0.05 was considered statistically significant. The same test compared the total number of CD3+ T cells and the percentage of p210-specific CD8+ T cells in morphologically stable versus unstable plaques. Population characteristics and pentamer-positive cell data are presented as percentages.

## RESULTS

### POPULATION CHARACTERISTICS

We enrolled 49 consecutive patients, with 44,8% testing positive for HLA-A*02:01 (22 patients).

The characteristics of the HLA-A*02:01+ population are shown in Table 1. Of the 22 patients, 16 were males. The mean age of participants was 73,88,4 years old. Hypertension, diabetes, and family history of cardiovascular disease were present in 86,9%, 43,4%, and 52,1%, respectively. Dyslipidemia occurred in 82,6% of patients, and 73,9% of them were already on therapy with lipid-lowering drugs. The mean LDL and HDL cholesterol levels in HLA-A*02:01 patients were 85 mg/dL and 32 mg/dL, respectively, while the mean BMI was 26.12 kg/m^2^. Smokers made up 69.5% of the entire population. Only 21.7% had reduced renal function (eGFR < 60 ml/min); the mean percentage of carotid stenosis before CEA was 79.0 ± 0.08%. Regarding clinical presentation, 5 of the 22 patients developed symptoms-transient ischemic attack or minor stroke-in the previous 6 months before CEA.

**Table 1.** HLA-A*02:01+ patients’ clinical characteristics.

| **Variables** | **Patients**<br>**(22)** |
| --- | --- |
| **Clinical Characteristics** |  |
| Age (years ± SD) | 73,8±8,4 |
| Gender |  |
| Male [n, (%)] | 16 (72,7) |
| Female [n, (%)] | 6 (27,3) |
| Symptoms [n, (%)] | 5 (22,7) |
| Diabetes Mellitus [n, (%)] | 10 (43,4) |
| Familial history of CAD [n, (%)] | 12 (52,1) |
| Hypertension [n, (%)] | 20 (86,9) |
| Dyslipidemia [n, (%)] | 19 (82,6) |
| Smokers [n, (%)] | 16 (69,5) |
| eGFR < 60 ml/h [n, (%)] | 5 (21,7) |
| BMI [mean ± SD] | 26,1±4,3 |
| LDL levels [mean mg/dl ± SD] | 85±33 |
| HDL levels [mean mg/dl ± SD] | 43±13 |
| Use of lipid lowering therapies [n, (%)] | 17 (73,9) |
| % of carotid stenosis [average % ± SD] | 79±0,1 |

### P210 SPECIFIC CD8+ T CELL POPULATION IN HUMAN PLAQUES

Atherosclerotic plaques from patients who tested positive for HLA-A*02:01 showed a mean of 6,715.2 × 10^3 HPF CD3+ cells, representing the entire T cell population. Co-staining with the p210/A*02:01 Pentamer identified a mean of 3.42 × 10^3 HPF p210-specific CD8+ T cells, which was significantly different from controls (p<0.0001). Spatial analysis of stained A*02:01 (+) atherosclerotic plaques revealed that p210-specific CD8+ T cells are mainly located in the shoulders of the atheroma, with low fluorescent signal from the core areas (Fig. 2). To confirm the T cell phenotype, a second staining with CD8 antibody (green) was performed (Fig. 3). Symptomatic patients showed a higher inflammatory infiltrate of the plaque than asymptomatic patients (9,249 vs. 5,83 CD3+ cells). A trend was observed in symptomatic patients, who had a slightly higher number of p210-specific CD8+ T cells compared to asymptomatic patients (77% vs. 57%). However, this difference did not reach statistical significance (p = 0.0659), and there was a significant temporal bias between the symptom onset and plaque collection in this group of patients (4 to 90 days), suggesting a different developmental stage of plaque at the time of staining. Histopathological evaluation classified 36% of the samples as unstable (8 out of 22) and 64% as stable (12 out of 22). Unstable plaques were mainly associated with calcific nodules (4 out of 8), followed by 3 cases of rupture and 1 case of thrombotically active plaque. Stable plaques were primarily classified as fibrocalcific.

**Fig. 2.**
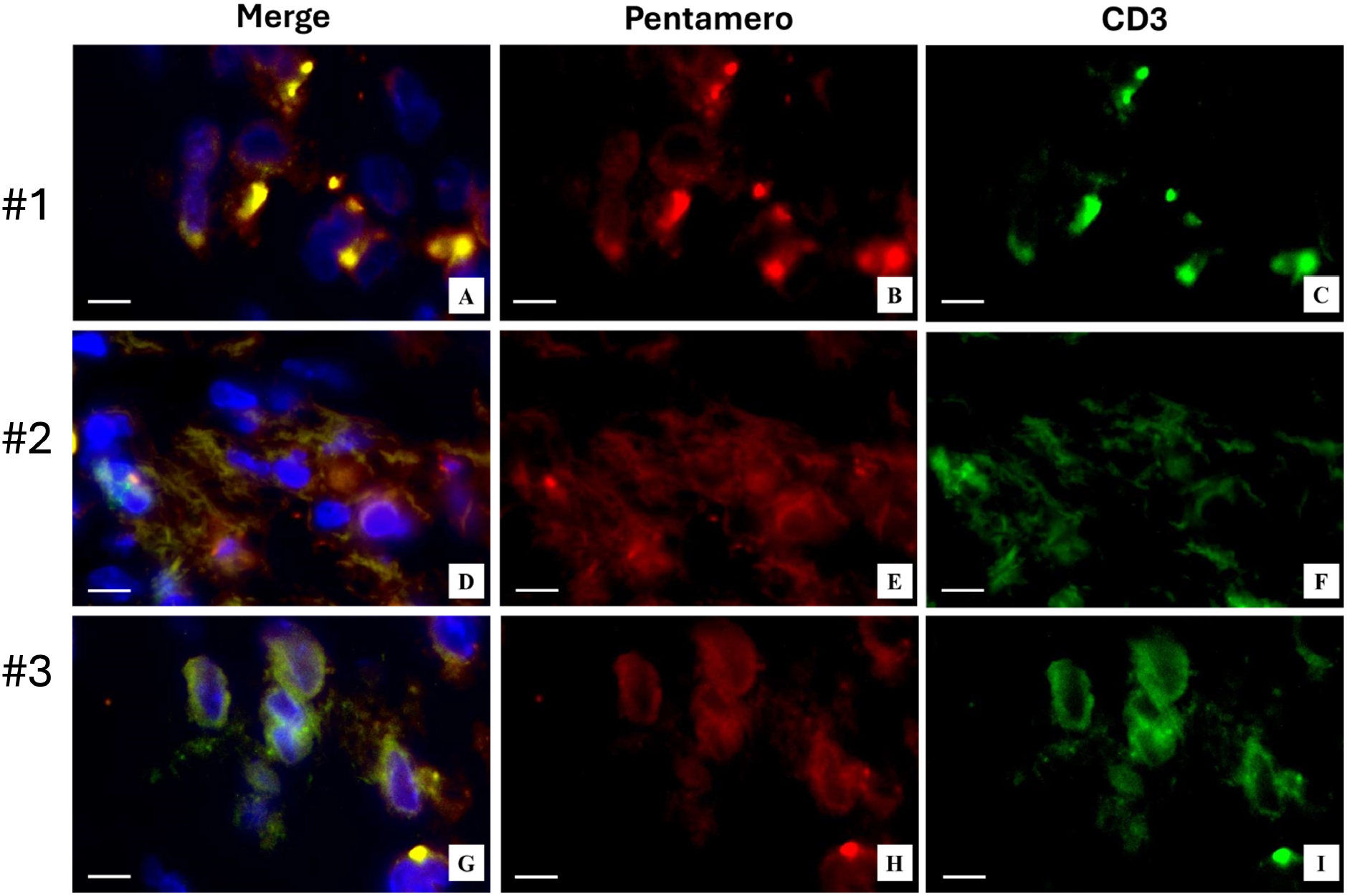
Immunohistochemistry of three symptomatic HLA-A*02:01+ patients (100X MAG).

**Fig. 3.**
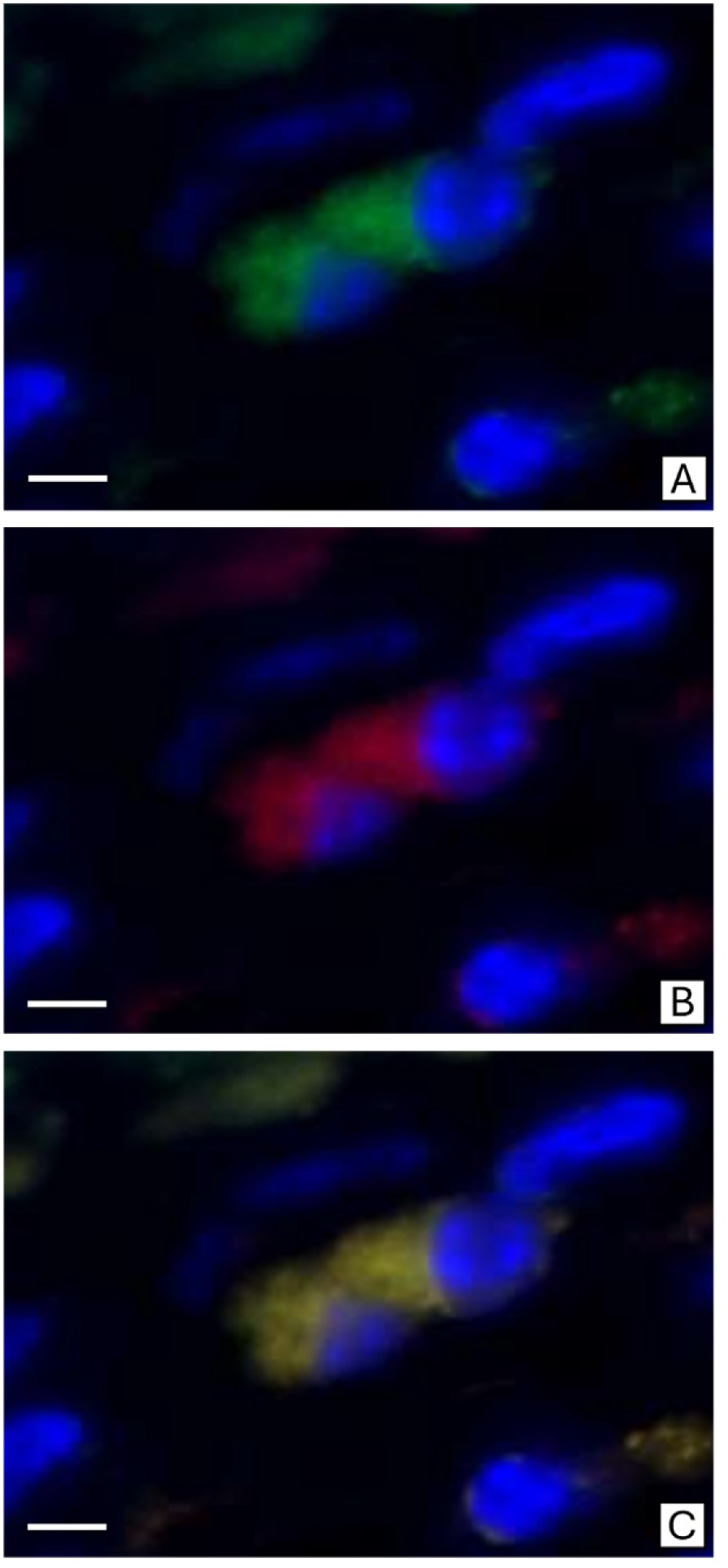
Pentamer-CD8 Dual Color Immunofluorescence. (A) The image shows CD8-positive cells (green) in a stable carotid plaque. (B) The image displays Pentamer-positive cells (red) in a stable carotid plaque. (C) The merged image of panels A and B demonstrates the co-expression of Pentamer and CD8 in inflammatory cells.

**Fig. 4.**
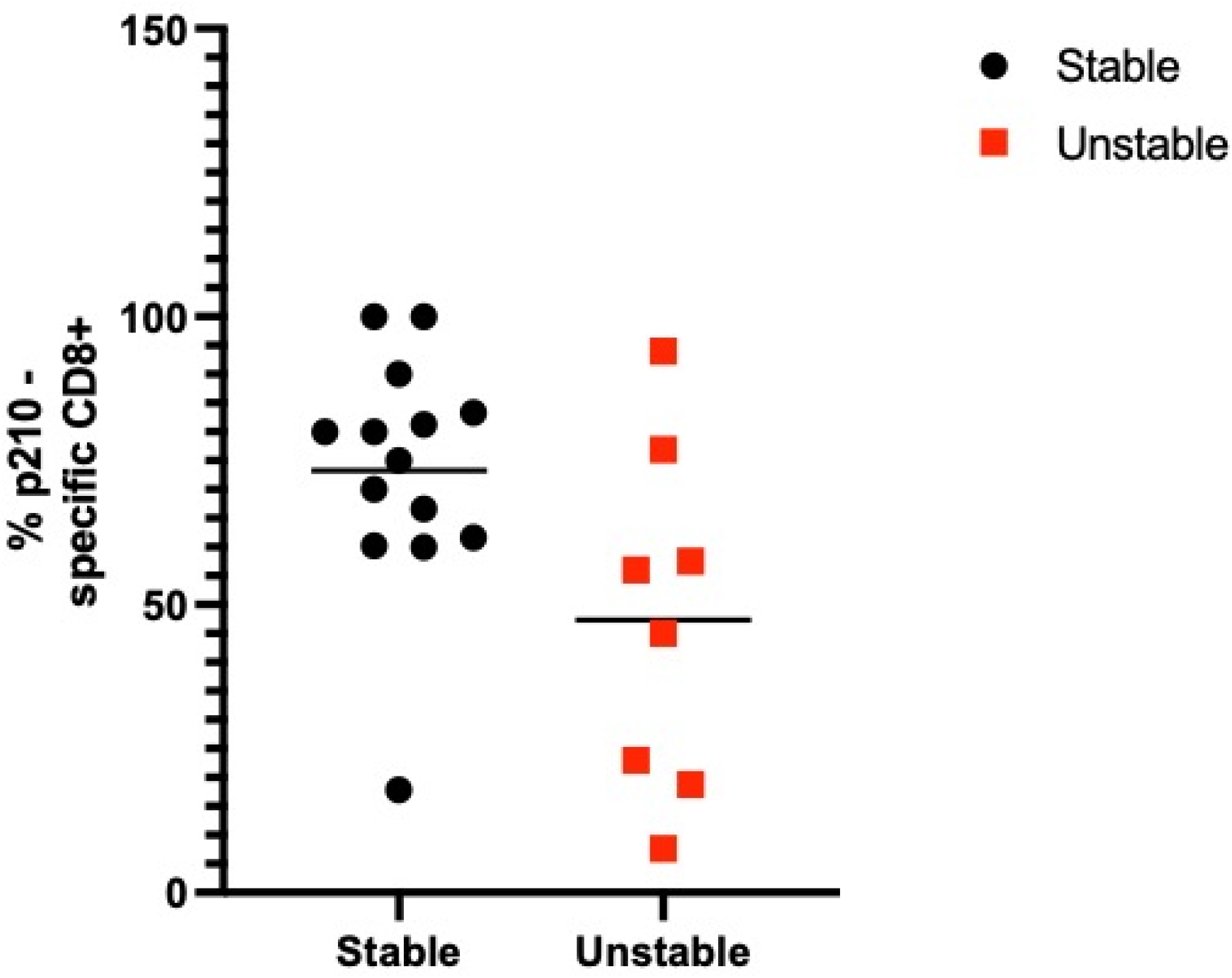
Difference in % of p210-specific CD8+ T cells in Stable and Unstable plaque. Median Stable 77,5. Median Unstable 50,5. P=0,027.

Interestingly, unstable plaques showed a lower number of p210-specific CD8+ T cells compared to stable plaques, as shown in Table 1 (p = 0.027 by a Mann-Whitney test).

## DISCUSSION

In recent years, identifying a self-antigen involved in human atherosclerosis has become “the missing link in the chain” for developing a potential vaccine. The present study is, to date, the first to examine the presence of a p210-specific CD8+ T cell population in HLA-A*02:01+ human atherosclerotic plaques.

The antigen receptors of CD4+ and CD8+ T cells are specific for peptide antigens presented by the HLA molecules. Regarding the HLA restriction phenomenon, a single T cell can recognize a specific peptide only when it is displayed by one of the many different HLA molecules that exist. Specifically, CD8+ T cells bind to class I HLA molecules and, once activated and differentiated into CD8+ cytotoxic T lymphocytes, perform their function.

In a previous study, Chyu et al. demonstrated the presence of p210-specific T cell responses in humans with atherosclerotic cardiovascular disease. They created a humanized atherosclerosis mouse model with an HLA-A*02:01/Kb chimera on an ApoE^−/−^ background. HLA-A*02:01 occurs with the highest frequency among Caucasians. ProImmune tested the p210 epitope binding to HLA*02:01 using the REVEAL assay, thus generating an ApoBKTTKQSFDL pentamer. By using such a pentamer, it has been possible to detect a small but significant population of P210-specific CD8+ T cells in peripheral blood mononuclear cells (PBMCs) from healthy HLA-A*02:01+ individuals^19^.

We used the same fluorochrome-labeled ApoBKTTKQSFDL pentamer to identify a p210-specific CD8+ T cell population in HLA*02:01+ human atherosclerotic plaques. All plaques from HLA*02:01+ patients showed varying numbers of p210-specific CD8+ T cells, mainly located in the shoulders of the atheroma.

In a previous study, our group demonstrated a strong association between hypertension, low HDL cholesterol (HDL-C), and a ratio of total to HDL-C greater than five with vulnerable and thrombotic carotid plaques. Additionally, hypertension, hypercholesterolemia, and low HDL-C levels were significantly linked to a high inflammatory infiltrate in the plaque. Consistent with these findings, all symptomatic patients in this study were hypertensive; four out of five were dyslipidemic and had lower HDL-C levels compared to asymptomatic patients (39. 39.25 mg/dL vs. 44. 7 mg/dL). Inflammatory infiltrates were more prominent in atherosclerotic plaques from symptomatic patients than from asymptomatic ones. Interestingly, symptomatic patients with TIA or minor stroke showed a higher percentage of p 210-specific CD 8 + T cells compared to asymptomatic patients. In a preclinical study, ApoE^-/-mice^ fed an atherogenic diet exhibited an increase in self-reactive p 210-specific CD 8 + T cell populations^18^, likely due to increased antigen exposure. Plaque rupture is preceded by a rise in effector memory T cells, followed by a decrease in total T cells. After plaque rupture in healed plaques, the decline in T cells is more pronounced for CD 4 + cells than for CD 8 + cells^23^. The finding that plaques from symptomatic patients contained higher amounts of CD 3 + T cells and a higher percentage of p 210-specific CD 8 + T cells compared to asymptomatic patients led us to hypothesize that continuous p 210 exposure activates the immune system, promoting the accumulation of these cells in the atherosclerotic plaque and contributing to its growth until complication (ulceration, rupture). However, a limitation of this study is the significant time gap between the acute cardiovascular event (ACE) and carotid endarterectomy (CEA) in the symptomatic subgroup. These patients underwent surgery within six months of an ACE (4 to 90 days), which is a wide timeframe to determine the precise stage of plaque evolution. Nonetheless, histopathological analysis revealed a reduced number of p 210-specific CD8+ T cells in unstable plaques (including calcific nodules, ruptures, and thrombotic plaques), suggesting a role for this cell population in plaque vulnerability. This aligns with a study by Van Dijk RA et al., which observed an increase in CD 8 + T cells in stable plaques (early fibroatheroma) compared to unstable plaques. Therefore, the presence of p 210-specific CD 8 + T cells could serve as a marker of plaque stability. Furthermore, in a mouse model, a protective effect of p 210-specific CD 8 + T cells has also been demonstrated.

Our findings offer new insights into the development of atherosclerosis and reveal a specific CD8+ T cell population targeting a self-antigen in human plaques, emphasizing the autoimmune origin of this silent killer.

Further studies are needed to characterize better the function and phenotype of the p210-specific CD8+ T cell population.

## CONCLUSION

This study, for the first time, identified a p210-specific CD8+ T cell population in human carotid atherosclerotic plaque from HLA-A*02:01+ patients, suggesting a potential role of autoimmunity in human atherosclerosis development. The presence of this cell population confirms p210 as a self-antigen in HLA-A*02:01+ patients. The translational significance of preclinical studies, combined with our findings, supports the potential effectiveness of p210 immunization in HLA-A*02:01+ individuals to reduce atherosclerosis.

## Notes

### Competing Interest Statement

The authors have declared no competing interest.

